# Comparative whole plastome and low copy number phylogenetics of the core Saccharinae and Sorghinae

**DOI:** 10.1101/2022.01.04.474895

**Authors:** Dyfed Lloyd Evans, Ben Hughes, Shailesh Vinay Joshi

## Abstract

Despite over 60 years’ worth of taxonomic efforts, the relationships between sugarcane (*Saccharum* hybrid cultivars), *Sorghum* and their closest evolutionary relatives remain largely unresolved. Even relationships between generally accepted genera such as *Miscanthus* and *Saccharum* have not been examined in any large-scale molecular detail. Genera such as *Erianthus*, *Miscanthidium* and *Narenga* pose even greater taxonomic contention. *Erianthus* is not monophyletic and *Erianthus* sect. *Ripidium* (Valdés and Scholz 2006, Lloyd Evans et al. 2019a; Welker et al. 2019) represents a distinct and separate genus, *Tripidium* Scholz. *Miscanthidium* is placed within *Miscanthus* by many workers, whilst the New World *Erianthus* species and *Narenga* are currently placed within *Saccharum*. As these species represent a significant portion of the gene pool that sugarcane breeders use for introgression into sugarcane, their taxonomic placement and relationships to *Saccharum* are of significant economic import. *Erianthus* species from the Americas have not been significantly employed in sugarcane breeding and may represent an untapped genetic resource. In an attempt to resolve the taxonomic relationships of these genera, we have assembled three novel chloroplasts, from *Miscanthidium capense*, *Miscanthidium junceum* and *Narenga porphyrocoma* (this latter assembled from transcriptomic and long read data). In parallel, five low copy number loci have been assembled from species within *Saccharum*, *Miscanthus*, *Sarga* and *Sorghum*. Phylogenetic analyses were performed using both low copy number genes and whole chloroplasts. The phylogenetic results were compared with karyotype data to circumscribe the genera most closely related to sugarcane. We reveal that genera *Miscanthus* and *Saccharum* are monophyletic and have never undergone polyploidization outside their own genera. Genera *Erianthus*, *Miscanthidium* and *Narenga* are allopolyploids, which excludes them from being members of *Saccharum* and *Miscanthus*. Moreover, all three of these genera have divergent evolutionary histories. We therefore support the use of the genera *Miscanthus*, *Miscanthidium*, *Erianthus* (for the New World Species) and *Narenga* for those species and genera most closely allied to Saccharum. Our data demonstrate that all these genera should be excluded from *Saccharum sensu lato*.

---

Sugarcane (complex hybrids of *Saccharum cultum* Lloyd Evans and Joshi, *Saccharum officinarum* L. and *Saccharum spontaneum* L. [Lloyd Evans and Joshi (2016)]) ranks amongst the top-ten crop species worldwide. Sugarcane also provides between 60 and 70% of total world sugar output and is a major source of bioethanol (Reddy et al. 2008). *Saccharum officinarum* is the type species for genus *Saccharum* L. *Saccharum sensu stricto* (*s.s.*) consists of four true members of genus saccharum: *Saccharum spontaneum*, *Saccharum robustum*, *Saccharum officinarum* and *Saccharum cultum* (Lloyd Evans and Joshi, 2016). However, many authorities also treat *Saccharum* in a broader sense (*Saccharum sensu lato*). For example, Kew’s GrassBase currently recognizes 36 species within *Saccharum* (http://www.kew.org/data/grasses-db/sppindex.htm#S).

Mukherjee (1957) first defined the ‘saccharum complex’ as group of potentially interbreeding genera believed to be involved in the origins of modern sugarcane hybrids (*Saccharum* ×*officinarum*/*Saccharum* ×*cultum* [Lloyd Evans and Joshi (2016)]). This concept was further refined by Clayton and Renvoize (1986) who extended the subtribe Saccharinae to include the genera: Erianthus Michaux, Eriochrysis P. Beauv., Eulalia Knuth, Eulaliopsis Honda, Homozeugos Stapf, Imperata Crillo, Lophopogon Hack, Microstegium Nees, Miscanthus Andersson, Pogonatherum P. Beauv., Polliniopsis Hayata, Polytrias Hack, Saccharum and Spodiopogon Trin. (as a result many of these genera have been re-classified as *Saccharum* and genus *Saccharum* now comprises between 35 and 40 species, mostly from the tropics and sub-tropics). The suggestion being, that all these genera are closely allied to Saccharum and were actually involved in the evolution of sugarcane’s ancestors. This paper has had considerable taxonomic influence and, for example, both the New World and Old World gernera of *Erianthus* as well as *Narenga porphyrocoma* are now all included within *Saccharum sensu lato*.

However, some authorities have considered *Erianthus* as a distinct genus (containing 28 species) originating in the Americas, Africa, Europe and Asia (Mukherjee, 1958; Molina, 1981; Watson and Dallwitz, 1992). Morphologically, *Erianthus* is distinguished from *Saccharum* by the presence of awned spikelets in the former and awnless spikelets in the latter (Mukherjee, 1958). However, the Old World species of *Erianthus* are grouped in a different section (*Erianthus* sect. *Ripidium* (Trin.) Henrard) or a distinct genus (*Tripidium* H. Scholz) by various authors (Grassl, 1972; Besse et al., 1997; Hodkinson et al., 2002; Valdés and Scholz, 2006). *Narenga* Bor and *Miscanthidium* Stapf. are also accepted as distinct genera by a few authors (e.g., Clayton, 1972; Watson and Dallwitz, 1992; Hodkinson et al. 2013) rather than including their species in *Saccharum* (*Tripidium*) or *Miscanthus* (*Miscanthidium*).

The circumscription of *Saccharum* remains highly controversial and has changed significantly over the past century. Several phenetic studies have indicated strong molecular differentiation between *Saccharum* and *Erianthus* (Besse et al., 1998; Nair et al., 2005; Selvi et al., 2006). Conversely, a phylogenetic analysis based on the internal transcribed spacer (ITS) of the nuclear ribosomal DNA (Hodkinson et al., 2002) found no support for this division, even though this study suggested that *Saccharum s.l*. is polyphyletic. Even the taxonomic delimitation between *Saccharum* and *Miscanthus* is not clear, with intergeneric hybrids occurring between them (Clayton and Renvoize, 1986; Hodkinson et al., 2002). However, a recent paper (Lloyd Evans and Joshi, 2016) comparing whole plastid genomes revealed a clear phylogenetic separation of *Miscanthus* and *Saccharum*, with the genera diverging at least 3.4 million years ago.

Previous studies (Estep et al. 2014, Welker et al. 2015) using low copy number genes and Soreng et al. (2015) using whole chloroplast analyses, demonstrated that the subtribe Saccharinae Griseb. was not monophyletic. However, both these studies had incomplete samplings of Saccharum s.s., as well as sugarcane’s known close relatives (*Miscanthus*, *Miscanthidium*).

As the Saccharum complex/Saccharinae comprises the gene pool that sugarcane breeders use when attempting to introgress useful characteristics into sugarcane the true relationship of these genera and species to each other, as determined by molecular techniques is of considerable import and relevance. This is especially the case, as modern molecular techniques do not support the concept of a ‘saccharum complex’ (D’Hont et al. 2008). Moreover, there is increasing evidence that *Saccharum* is a well-defined lineage that diverged over a long evolutionary period from the lineages leading to the New World *Erianthus* and Old World *Miscanthus* genera (Grivet et al. 2004; Welker et al. 2015; Lloyd Evans and Joshi, 2016).

The recent finding (Shi et al. 2016) that plant chloroplasts are completely transcribed means that it is theoretically possible to assemble whole plant chloroplasts from transcriptomic data. This means that we can begin to fill in the gaps in assembled Saccharinae chloroplast data by mining transcriptomic datasets.

One of the problem species within Saccharum has always been *Saccharum narenga* (Nees ex Steud.) Wall. ex Hack. In 1940 Bor moved the species from *Saccharum* into *Narenga* as *Narenga porphyrocoma* (Hance ex Trimen) Bor (based on the basionym *Eriochrysis narenga* Nees ex Steud.). Bor excluded this species from genus *Saccharum* due to its possession of the following anomalous characters: non-coriaceous glumes, the absence of non-flowering stems, the plant generally being less ‘robust’, a reduced panicle, the presence of secondary branches only on the lowest lateral branches of the panicle and the presence of a fourth glume in the florets (Janaki-Ammal, 1942). To add to the confusion, the NCBI’s taxonomy (https://www.ncbi.nlm.nih.gov/taxonomy/?term=saccharum%20narenga) defines this species as *Narenga porphyrocoma*, whilst Kew’s GrassBase defines it as *Saccharum narenga* (http://www.kew.org/data/grasses-db/www/imp09051.htm). Tropicos also defines the accepted name as *Saccharum narenga* (http://www.tropicos.org/Name/25509769? tab=acceptednames).

Though *Saccharum narenga* (*Narenga porphyrocoma*) has not been used extensively in sugarcane introgression breeding, it does possess a number of characters that would be beneficial to sugarcane: broad-range disease resistance, pest resistance, root pest resistance, drought resistance and waterlogging resistance, early flowering and high tillering (Heinz, 2015; Shrivastva and Srivastva 2016).

*Saccharum narenga* has a natural range that extends from India through Burma and Thailand to south-eastern China and Taiwan, with a rump population in subtropical Ethiopia (Clayton et al. 2006). It is classed as ‘potentially weedy’ (Cheavegatti-Gianotto et al. 2011). As such, its taxonomic relationships to *Saccharum s.s*. is of great import as it could be a bridge species, allowing transgenes to escape from sugarcane into the wild.

Following Estep et al. (2015) we have assembled five low copy number gene regions (*Aberrant panicle organization1* (*apo1*), *Dwarf8* (*d8*), two exons (7 and 8) of *Erect panicle 2* (*ep2*), and *Retarded palea 1* (*rep1*)) from four *Miscanthus* accessions, one *Miscanthidium* accession and four *Saccharum* accessions along with *Sorghum propinquum* (Kunth) Hitchc. and *Sarga versicolor* (Andersson) Spangler. In addition, the chloroplast genome of *Saccharum narenga* was assembled from high depth transcriptomic data.

Comparison of plastome and low copy number genomic data indicate that Old World *Erianthus* species and *Miscanthidium* species are hybrids of a common ancestral lineage, sister to both *Saccharum* and *Miscanthus* with an early-diverging species that is sister to all the crown *Saccharum* species. *Saccharum narenga* has the same ancestry, but about three million years ago it underwent a back-hybridization with a different early-diverging *Saccharum* sister. The lineages leading to *Miscanthus sinensis* Andersson and *Miscanthus sacchariflorus* (Maxim.) Benth. & Hook. f. ex Franch. reciprocally hybridized with each other, leading to a duplication in the base genome. Saccharum emerges as being unusual in the entire grouping, not having undergone any ancestral hybridizations outside its own genus subsequent to divergence from *Miscanthus*.

We conclude that *Saccharum*, *Erianthus*, *Miscanthidium*, *Miscanthus*, *Tripidium* (*Erianthus* sect. *Ripidium*) and *Narenga* are separate genera with distinct evolutionary histories. However, an ancestral member of *Saccharum* was involved in the origins of *Erianthus* and *Miscanthidium*. Both chloroplast and genomic data supports inclusion of genus *Sarga* within the saccharinae subtribe, but more studies are required to ascertain the placement of *Sorghum*. Though *Narenga porphyrocoma* is genetically closer to *Saccharum* (at both the plastome and gross genomic level) than any of the other genera it should be excluded from genus *Saccharum* on the basis of different physical characteristics, different evolutionary ancestry and different base chromosomal counts.

## Materials and Methods

### Plant Materials and Transcript/Chloroplast Extraction

Leaf material for Saccharum narenga (Narenga porphyrocoma) was collected in Malaysia (isolatedd to the north of Kak Bukit), voucher Mah:CSS:013. Chloroplasts were captured with conserved tRNA magnetic bead bound primers and sequenced with ONT MinION as described previously (Lloyd Evans et al. 2019b).

### Low Copy Number Locus Assembly

Low copy number regions were assembled for the following genes: *Aberrant panicle organization1* (*apo1*), *Dwarf8* (*d8*), two exons (7 and 8) of *Erect panicle* 2 (*ep2*), and *Retarded palea 1* (*rep1*). We assembled these regions from the following GenBank sequence read archive (SRA) accessions: *Sorghum propinquum* (SRR072055); *Sarga versicolor* (SRR427176); *Miscanthidium junceum* (SRR396848 and SRR396849); *Miscanthus sacchariflorus* var. Hercules (SRR486748); *Miscanthus x giganteus* (SRR407328 and SRR407325); *Miscanthus transmorrisonensis* (SRR396850); *Saccharum* hybrid LCP85-385 (SRR427145); *Saccharum* hybrid RB72454 (SRR922219) and *Saccharum spontaneum* SES234B (SRR486146). Narenga porphyrocoma transcripts were assembled based on the Miscanthidium junceum assemblies using the SRA accessions (<u>SRR3399436</u> and <u>SRR6322472</u>) from these assemblies capture primers were developed (Supplementary Appendix 1) and these were employed for magnetic bead capture and MinION sequencing (Lloyd Evans 2021). *Miscanthus sinensis* cv. Andante genomic reads were kindly donated by CSS, Waterbeach, Cambridge, UK. Gene loci were assembled as described previously (Lloyd Evans and Joshi, 2017) using the closest orthologue for each gene for Mirabaiting and SPAdes for assembly. Where a second copy of the gene co-assembled (typically as a shorter, second, assembly contig), this was co-assembled along with the primary gene copy. In *Miscanthus*, *M. sinense* was used to bait the A copy and *M. sacchariflorus* was used to bait the B copy for each species. For *Saccharum*, *Saccharum spontaneum* was used to bait the spontaneum copies of the gene from *Saccharum* hybrid LCP85-385. In addition, all species that were near neighbours of sugarcane were used to bait data from *Saccharum* hybrid LCP85-385 to ensure that no alternate copies of any gene were present in the genome. All *Miscanthus sinense* cv Andante assemblies were have been deposited in EMBL/GenBank under the project identifier PRJEB22229 and *Narenga* chlroplast assemblies were deposited in project ????????. Finally, Narenga transcript sequences were deposited in project ?????. All other gene assemblies were deposited in the Dryad repository along with their optimized alignment (DOI available upon acceptance).

### Chloroplast Assembly

The *Miscanthidium junceum* chloroplast was assembled from SRA datasets: SRR396848 and SRR396849, as described previously (Lloyd Evans and Joshi 2016) using *Saccharum spontaneum* and *Miscanthus sinensis* assemblies for baiting. The *Miscanthus* x *giganteus* plastome sequence was assembled from the SRA datasets SRR407328 and SRR407325 and the *Sarga versicolor* chloroplast was assembled from the SRA accession SRR427176, using the same methodology. The *Miscanthidium capense* Joshi1 chloroplast genome was assembled using 13 novel PCR primers (detailed in supplementary Appendix 2). Leaf samples were derived from a plant at the South African Sugarcane Research Institute living collection, Durban. Illumina short reads were derived from the PCR products and were assembled using SPAdes, as described previously (Lloyd Evans and Joshi, 2016) using the *Miscanthidium junceum* chloroplast as a reference.

The *Microstegium vimineum* chloroplast was assembled using a blend of RNA-seq data (NCBI SRA accession: ERR2040772) and previously assembled *Microstegium vimineum* chloroplast fragments (isolated from the NCBI using the query: “Microstegium vimineum”[organism] and (chloroplast or plastid)). The chloroplast fragments were converted to FASTQ with a custom PERL script. Reads corresponding to the chloroplast were isolated using mirabait with a k-mer of 27 employing the *Rottboellia cochinchinensis* and *Sorghastrum nutans* chloroplasts as a reference. All reads were co-assembled with SPAdes, using the *Rottboellia cochinchinensis* chloroplast as an untrusted reference. Eight contigs were obtained, which were scaffolded on the *Rottboellia cochinchinensis* chloroplast. Sequences corresponding to the gap regions (the largest of which was the entire IRa region) were excised from the *Rottboellia cochinchinensis* chloroplast and were surrounded by 1kbp regions from the *Microstegium vimineum* assembly. These isolated regions were baited against the read pool with mirabait and a k-mer of 25 before being assembled with SPAdes and stitched into the *Microstegium vimineum* assembly. After assembling and filling the gaps, followed by a second round of assembly to fill the remaining three small sequence gaps, the complete chloroplast sequence of *Microstegium viminium* was obtaind. The EMBL annotated version of the assembly can be downloaded from the Dryad digital repository.

Our previously assembled and annotated transcriptomic assembly of *Narenga porphyrocoma* (available from the Dryad digital repository) was also utilized in this study.

### Gene Region Alignments

Gene regions assembled in this study were merged with the equivalent gene regions from a subset of the data from Welker et al. 2015. All regions were aligned independently using SATÉ, with MAFFT as the aligner and Muscle as the sub-alignment joiner. Alignments were adjusted manually and trimmed. Each individual alignment was input into RAxML and a phylogenetic analysis (100 replicates) was run to identify the most likely tree. If alternate copies of genes were in the same position in the phylogeny these were linked. Genes were stitched together with a custom Perl script. If the positions of genes were uncertain, all alternate positions were generated. The phylogeny was run with RAxML and the sub-alignment giving the strongest phylogenetic signal was chosen. Using this methodology, we generated the optimal alignment (available from the Dryad repository). Chloroplast genomes were aligned, as previously described (Lloyd Evans and Joshi, 2016). A list of all chloroplasts used in the alignment, including GenBank accessions, voucher accession and publications are given in supplementary Appendix 3.

### Phylogenetic Analyses

The gene alignment was divided into the five constituent gene regions and models of DNA evolution were determined using PartitionFinder (Lanfear et al. 2014) and the AICc criterion. For each partition, the GTR + Г model was chosen. To determine the best topology, two independent runs of RAxML RAxML (version 8.1.17) (Stamakis, 2006), using different seeds were run, with 100 replicates. Both runs yielded the same best tree topology and this was used as the reference for all future analyses. Concatenated trees were reconstructed using both maximum likelihood (ML) and Bayesian approaches and rooted at *Arthraxon*. The ML tree was estimated with RAxML using the GTR + Γ model for all 5 partitions, and 6000 bootstrap replicates. The Bayesian tree was estimated using MrBayes v.3.2.1 (Ronquist and Huelsenbeck 2003) using a gamma model with six discrete categories and partitions unlinked. Two independent runs with 25 million generations each (each with four chains, three heated and one cold) were sampled every 1,000 generations. Convergence of the separate runs was verified using AWTY (Nylander et al. 2008). The first six million generations were discarded as burn-in. The ML trees and the MB trees were mapped onto the best topology from the initial RAxML run with SumTrees.

For the whole chloroplast alignment, the alignment was divided into LSC, IRA, SSC and IRB partitions. The GTR + Г partition was chosen for all partitions. Maximum likelihood and Mr Bayes runs were executed as described above for the low copy number gene data.

Divergence times on the whole chloroplast alignment were estimated using BEAST 2.4.4 (Drummond et al. 2012), on a 24-core server running Fedora 25, using four unlinked partitions (as above). The concatenated analysis was run for 50 million generations sampling every 1,000 under the GTR + Γ model with six gamma categories. The tree prior used the birth-death with incomplete sampling model (Drummond et al. 2012), with the starting tree being estimated using unweighted pair group method with arithmetic mean (UPGMA). The site model followed an uncorrelated lognormal relaxed clock (Drummond et al. 2006). The analysis was rooted to *Arundinella*, with the age of Zea mays divergence estimated as a normal distribution describing an age of 13.8 ± 2 million y (Estep et al. 2014). Convergence statistics were estimated using Tracer v.1.5 (Rambaut et al. 2013) after a burn-in of 15,000 sampled generations. Chain convergence was estimated to have been met when the effective sample size was greater than 200 for all statistics. Ultimately, 30,000 trees were used in SumTrees to produce the support values on the most likely tree (as determined above) and to determine the 95% highest posterior density (HPD) for each node. All final trees were drawn using FigTree v.1.4.0 (tree.bio.ed.ac.uk/software/figtree/) prior to finishing in Adobe Illustrator. Final alignments and trees are available from TreeBase (TB2:S21154 and TB2:S21155).

### Karyotyping

Images from the original publications (*Sarga vrsicolor*: Sun et al. (1994); *Miscanthus sinensis*: Cramiec-Głąbik et al. (2012); *Miscanthidium capense*: Strydom et al. (2000) and Hoshino and Davidse, (1988); *Erianthus giganteus*: Burner and Webster, (1994); *Narenga porphyrocoma*: Jagathesan and Sreenivasan (1967), *Saccharum spontaneum*: Ha et al. (1999); *Saccharum officinarum*: Li et al. (1959) and Divinagarcia and Ramirez, (1976)) were scanned in grayscale at 1200dpi prior to export as tiff files. Images were imported into Adobe Photoshop and were adjusted to improve contrast and sharpness. Images were re-exported in TIFF format and imported into the image analysis toolkit, icy (http://icy.bioimageanalysis.org). The Painting plugin (http://icy.bioimageanalysis.org/plugin/Painting) was employed to create a mask around each chromosome, with centromere to P-arm and centromere to Q-arm lengths determined by icy’s viewer pane tools. Three independent measurements were taken in each case and written to Excel files. A custom Perl script read in the file and determined chromosomal arm lengths, scaling the measurements based on the scale bar information from the original publications. The script output chromosome descriptor files in ‘.cfg’ format which were used as input to coloredChromosomes.pl (Böhringer et al. 2002) using custom configuration files for each species. The outputs (PostScript files) were imported into Adobe illustrator for integration prior to export as a single PDF file.

## Results

### Gene Assembly

Genes and gene fragments were assembled using a bait and build process, as described previously (Lloyd Evans and Joshi, 2017). For our 11 novel species/accessions (see Materials and Methods for details), all genes and gene fragments were assembled, except for the following: *apo1* — *Miscanthus transmorrisonensis* A, *Miscanthus sacchariflorus* A, *Saccharum* hybrid RB7234; *d8* — *Saccharum* hybrid RB7234, *Sorghum propinquum*, *Miscanthus sacchariflorus* B (partial); *rep1*: *Saccharum* hybrid RB7234. As a result, all species (including alternates) meet the 3/5 sequences present chosen as a minimum cutoff for phylogenetic studies established by Estep et al. (2014). Sequencing and assembly of the *Narenga porphyrocoma* transcripts yielded 100% identity with transcript based sequencing.

### Chloroplast Assembly

The two *Miscanthidium* chloroplasts were assembled as described previously (Lloyd Evans and Joshi, 2016). The *Miscanthidium junceum* chloroplast was 141 149 bp in length and the *Miscanthidium capense* chloroplast was 141 213 bp long (both had the same gene content). The same methodology was employed in the assembly of the Miscanthus x giganteus (141 220 bp long) and Sarga versicolor (140 951 bp long) chloroplasts. These latter two chloroplast assemblies had the same gene contnt and gene organization as the two *Miscanthidium* assemblies. Images of the four chloroplast assemblies generated in this study are shown schematically in Figure 1. The RNA-seq based assembly of *Narenga porphyrocoma* was 141 213 bp long.

**Figure 1.**
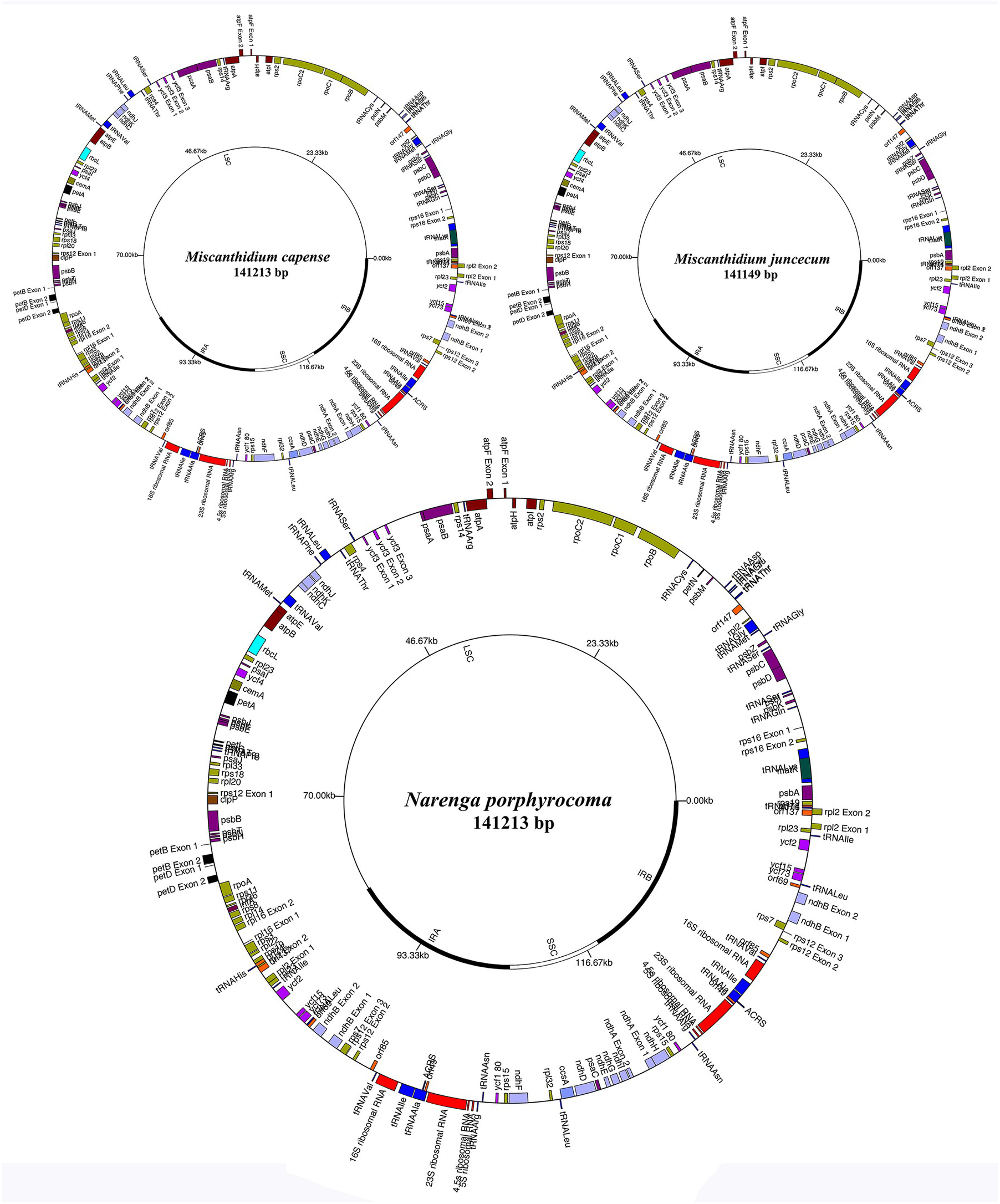
Schematic images for three complete chloroplast assemblies. Shown are three schematic drawings for the three chloroplast genomes assembled and annotated in the present study. Top left is the chloroplast of *Miscanthidium capense*, assembled from chloroplast PCR fragments. Top right is the chloroplast of *Miscanthidium junceum*, assembled from whole genome sort read data. Bottom is the chloroplast of *Narenga porphyrocoma*, assembled from transcriptomic data. Protein coding genes are shown on the outer track, with forward strand genes on the outside and reverse strand genes on the inside. The inner track shows the extent of the large single copy region (LSC), small single copy region (SSC) and the two inverted repeats (IRA and IRB). Images were drawn with GenomeVx prior to finishing with Adobe Illustrator.

### *Narenga* MinION Chloroplast Assembly

The MinION data for *Narenga porphyrocoma* was assembled with Canu (Koren et al. 2017) and the assembly graph was analyzed with unicycler (Wick et al. 2017) this revealed two distinct isoforms with the difference between them being an inversion of the SSC (short single copy region) with the ratios of the canonical and reversed forms being 52%:48%. The MinION based assemblies and RNA-seq based assemblies were 141 217 bp long, the lengh increased as compared with the transcript-based assembly due to a 4 bp insertion in the LSC. Apart from this, the sequences differed by 22 sequence alterations due to DNA to RNA transcription and 6 sequence alterations due to differing origins of the sequences.

### Low Copy Number Gene Phylogeny

Low copy number gene evolution for the Andropogoneae has been extensively analyzed previously (Estep et al. 2014; Welker et al. 2015). Indeed, the backbone of our phylogeny (Fig. 2) is based on data generated by these studies. However, we extend these studies by providing additional data for *Sarga versicolor, Miscanthus, Miscanthidium and Saccharum*. This fills in the gaps in the previous studies. The sister relationship of *Sarga* and *Saccharum s.l*. is well supported (85% ML, 1 BI) and monophyly of *Saccharum s.s*. is 100% supported. There is an early split between the ‘B’ genomes of *Miscanthidium* and *Erianthus* and this clade is also 100% supported. The next split is between a clade formed of *Pseudosorghum fasiculaire* and *Miscanthus* and the ‘A’ genomes of *Miscanthus*, *Erianthus*, *Narenga* and *Saccharum*. This split has poor ML support (62%) but good MB support (0.94) — allied with the short internal branches this is probably a sign of rapid radiation. The sister relationship of *Pseudosorghum* and *Miscanthus* has good support (83% ML, 0.9 MB) and monophyly of *Miscanthus* (as well as the split between the A and B genomes of *Miscanthus*) has 100% support. The split between the ‘A’ genome of *Miscanthidium*/*Erianthus* and *Saccharum s.s*. has good support (79% ML, 1 MB). In addition, the sister relationship of the *Narenga* ‘B’ genome and core *Sacharum* has decent support (65% ML, 0.99 MB). Monophyly of core *Saccharum* is 100% supported.

**Figure 2.**
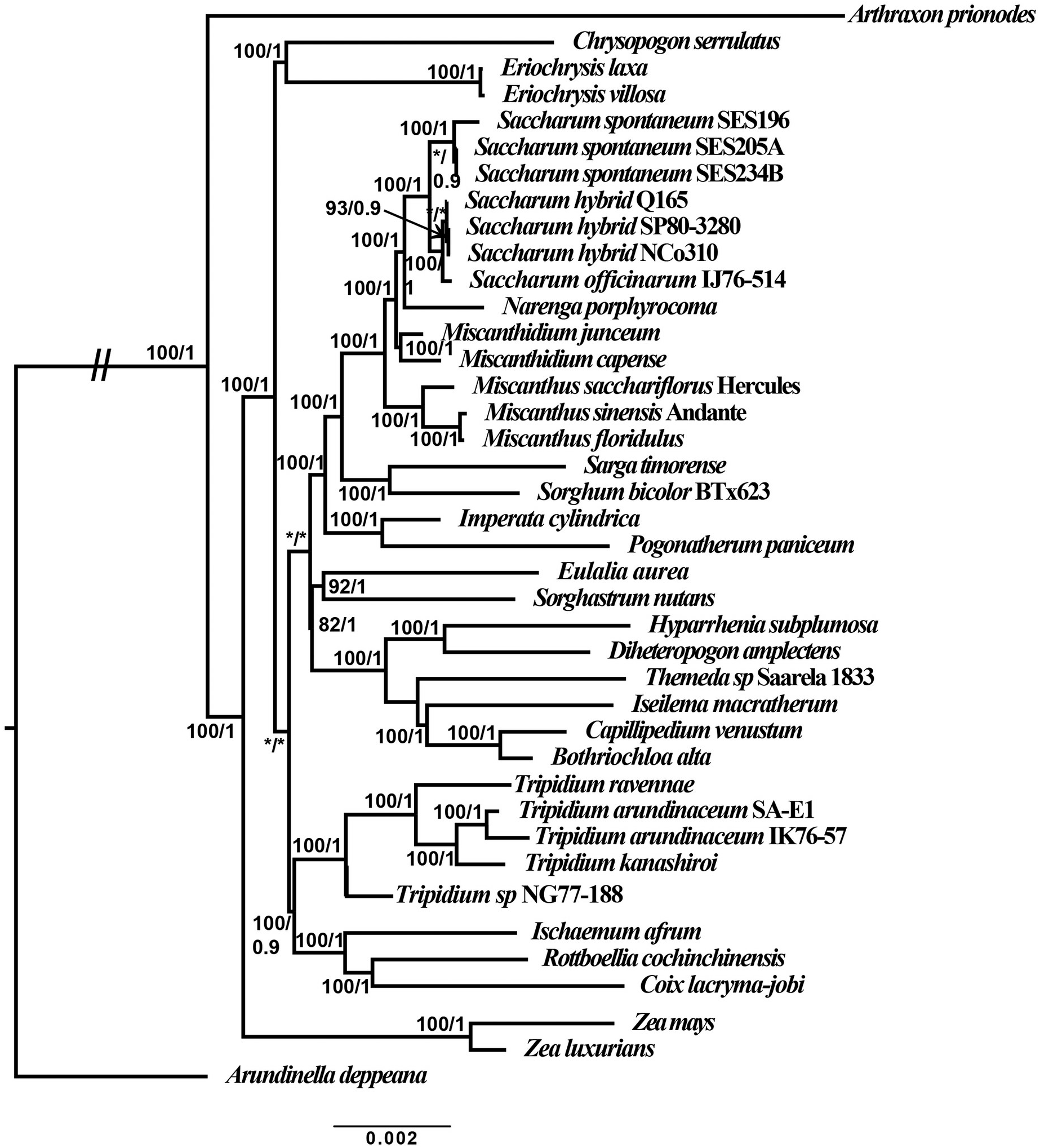
Whole Plastid Phylogram of Saccharum and Related Genera. Phylogram, representing the most likely topology from two independent runs of RAxML comparing whole chloroplast alignments of five *Tripidium* species along with the newly-assembled plastomes of *Miscanthidium junceum*, *Miscanthidium capense* and *Narenga porphyrocoma* with 31 representative members of the Andropogoneae. *Arundinella deppeana* is used as the outgroup. The two slashes (//) indicate that the long branch linking the outgroup to the remainder of the tree has been reduced by 50% so that internal relationships can be more clearly seen. Numbers next to nodes represent Maximum Parsimony bootstrap confidence/Bayesian inference where */* represents 100/1. The scale bar at the base of the phylogeny represents the expected number of substitutions per site. Images were drawn with FigTree prior to finishing with Adobe Illustrator. Currently accepted members of the Saccharinae are represented by a double dagger (‡) and currently accepted members of the Sorghinae are represented by an asterisk (*)

However, within *Saccharum*, branches have relatively poor support and the sister relationship of *Saccharum officinarum* and *Saccharum spontaneum* is not supported (this has 100% support in the chloroplast phylogeny), indicating that the dataset may have insufficient characters to resolve the relationships between close sister species (this is confirmed by ongoing studies with nine low copy number regions, D. Lloyd Evans, personal communication).

### Chloroplast Phylogeny

Chloroplast phylogeny, both phylogram (Figure 3) and chronogram (Figure 4) reveal that *Tripidum* (*Erianthus* sect *Ripidium*) is clearly distinct from *Saccharum s.s*. and *Saccharum s.l*. being 11.1 million years divergent. *Sorghum* and *Sarga* are sister to the core Saccharinae (6.6 million years divergent). *Miscanthus* diverged from Saccharum 3.7 Ma. *Miscanthidium* diverged from *Saccharum* 3 Ma and *Narenga* diverged from Saccharum 2.9 Ma. Within *Saccharum* we see the expected divergence of *Saccharum spontaneum*, *Saccharum officinarum* and the Sugarcane hybrid cultivars (based on *Saccharum cultum*). *Sorghum* is also clearly distinct from and 6.7 million years divergent from *Sarga*.

**Figure 3.**
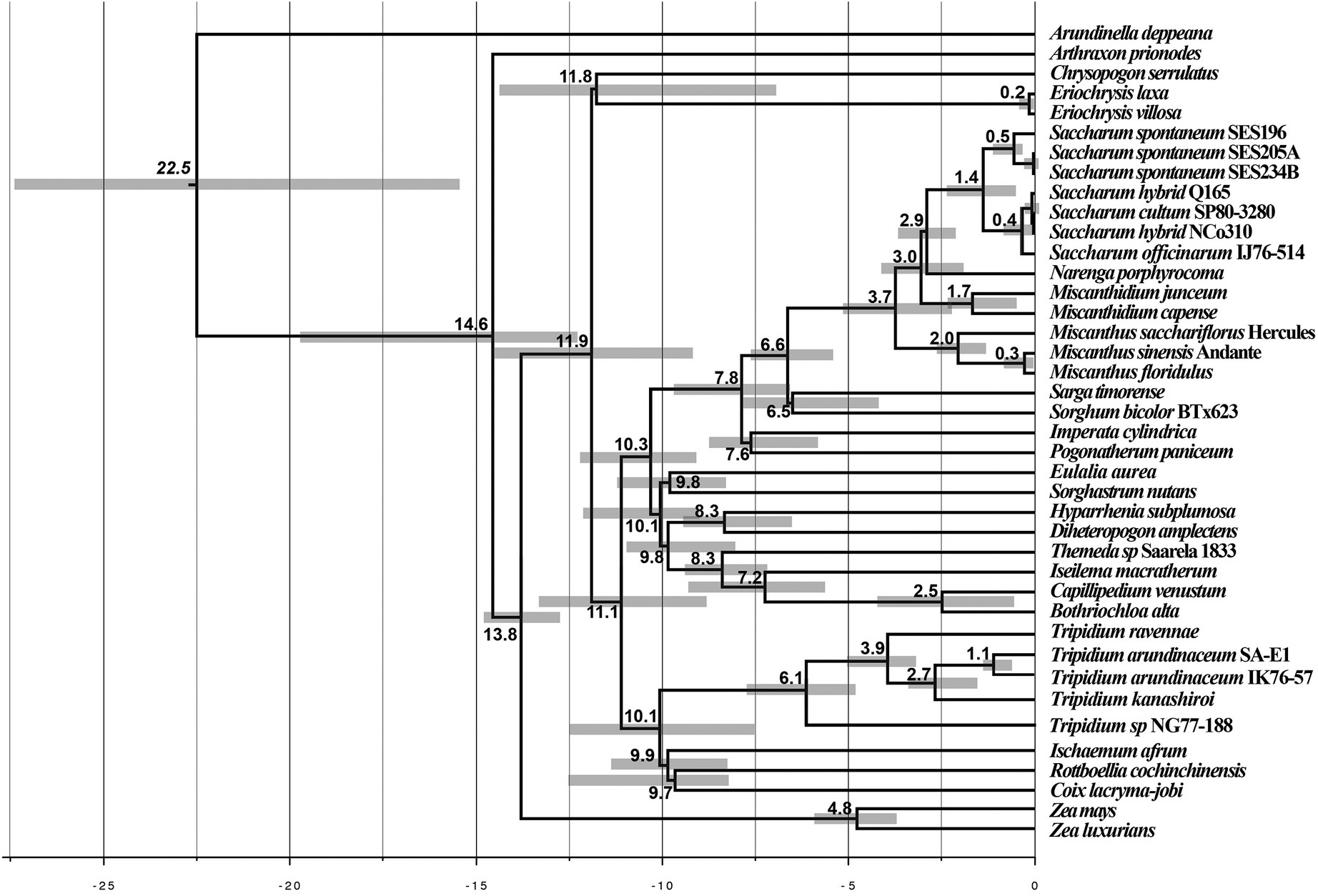
Chronogram for Andropogoneae divergence, focussing on the Saccharinae. The chronogram is based on whole plastid alignments showing the evolutionary distances (in millions of years) between the species and accessions analysed in this study. Numbers to the left of nodes show the age of the node in millions of years ago (Ma). Node bars give the 95% highest posterior density (HPD) for the node age. The scale at the bottom gives millions of years before present in 2.5 million year increments. *Arundinella deppeana* is employed as the outgroup. Images were drawn with FigTree prior to finishing with Adobe Illustrator.

**Figure 4.**
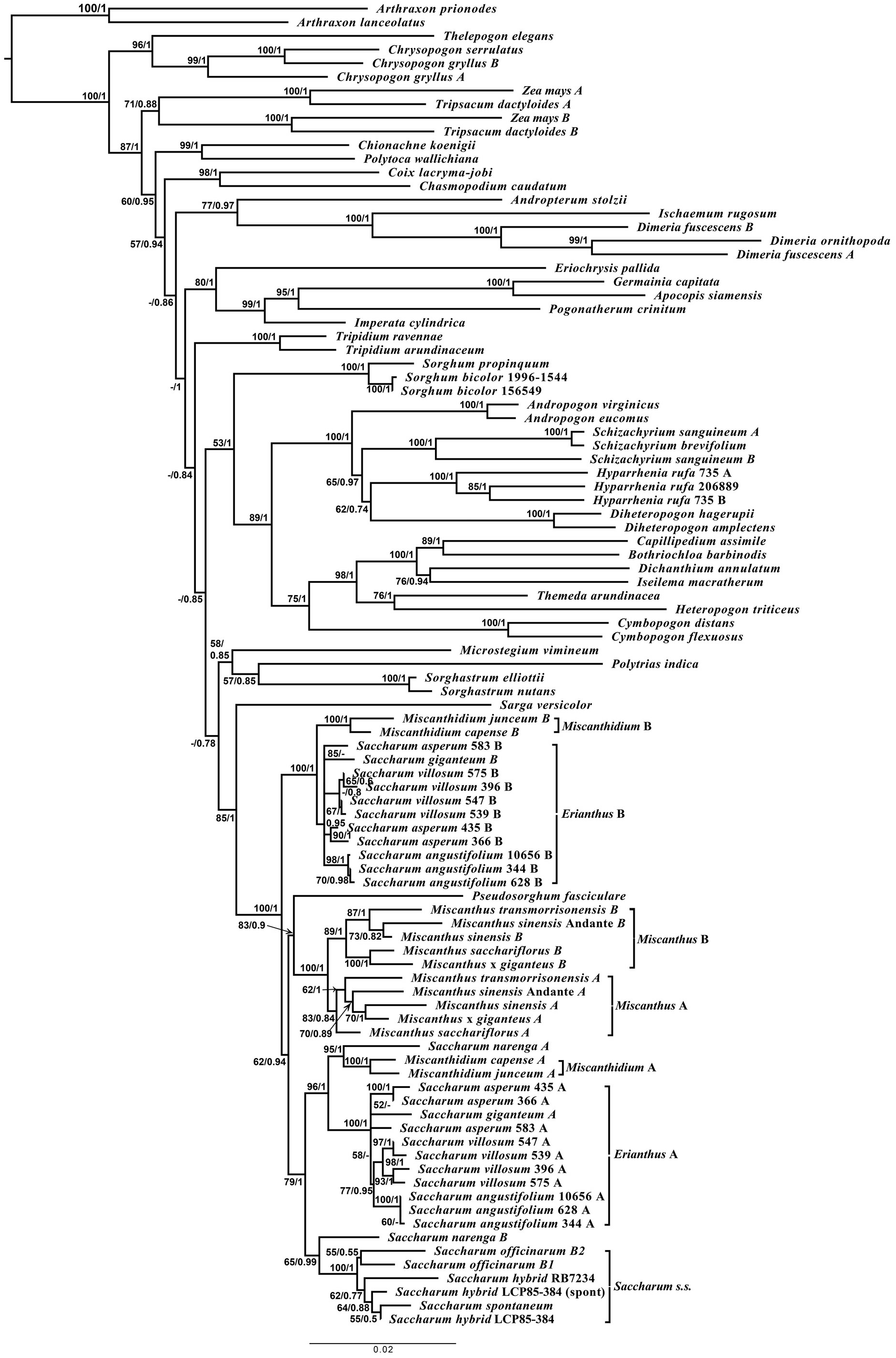
Phylogram based on five low copy number gene regions. A phylogram of Andropogoneae species, focussing on the core Saccharinae. Numbers next to nodes represent bootstrap confidence/Bayesian inference where ‘-’ represents confidences below the cutoff threshold (60% for MP and 0.7 for BI). The scale bar at the base of the phylogeny represents the expected number of substitutions per site. Divergent A and B genomes for *Erianthus*, *Miscanthidium*, *Miscanthus* and *Narenga* are shown by brackets. The phylogram was rooted on two *Arthraxon* accessions.

### Karyotype Analyses

By mining historical data we have managed to generate integrated ideograms depicting the basal (x) chromosome complement for *Sarga versicolor, Miscanthidium capense, Erianthus giganteus, Miscanthus sinensis, Narenga porphyrocoma, Saccharum spontaneum* and *Saccharum officinarum* (Fig. 5). In the majority of cases, chromosomes were small (on the order of 2– 3 µm in length), with a few anomalies in *Miscanthus sinensis*, *Saccharum officinarum* and, most notably, *Narenga porphyrocoma*. In all cases lengthened chromosomes or chromosomes with satellites were compatible with chromosomal fusion events. In the majority of cases, changes in base chromosomal counts were compatible with allopolyploid hybridization events.

**Figure 5.**
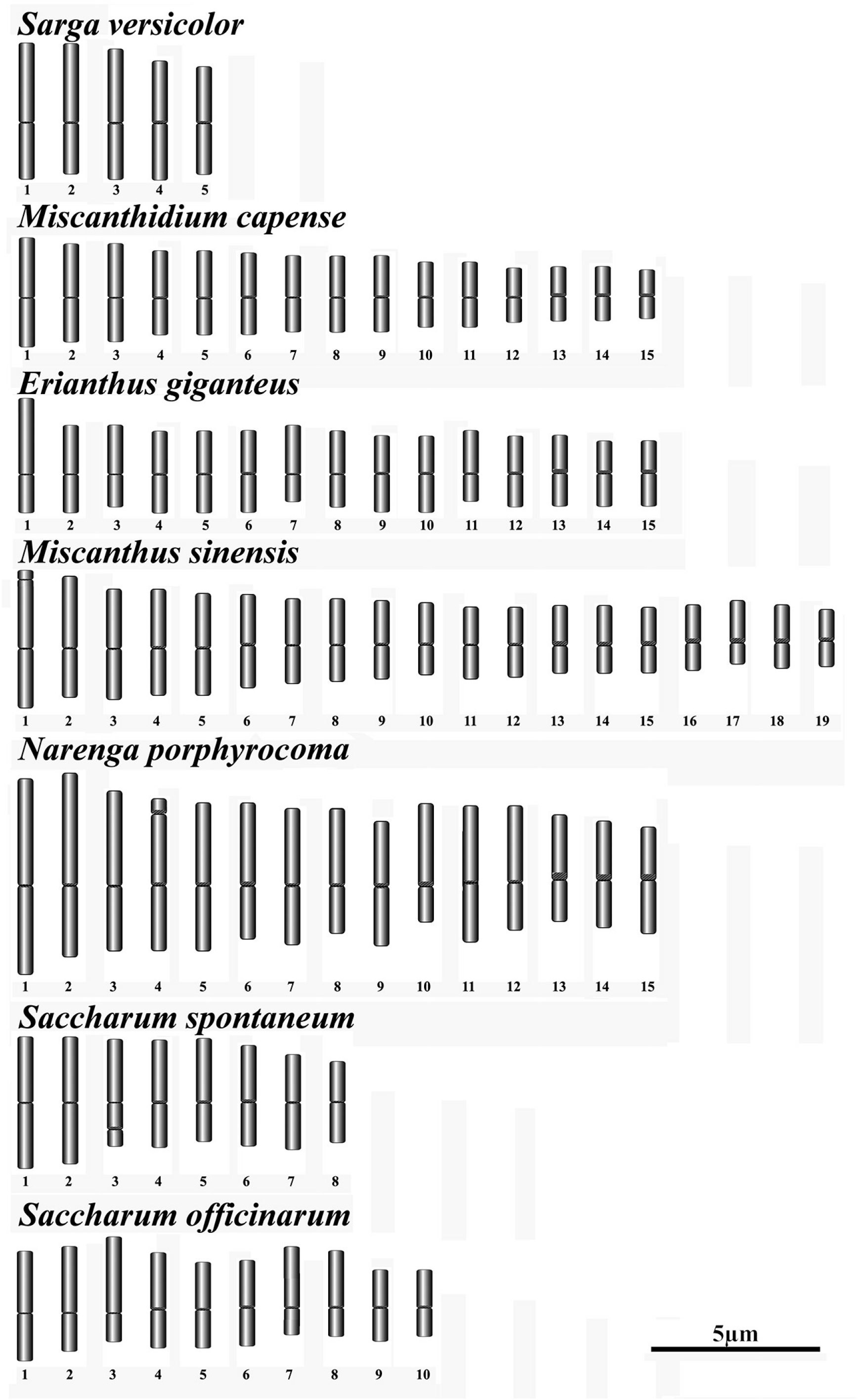
Schematic ideograms for members of the core Saccharinae. Ideograms were generated from previously published chromosomal analyses. P and Q arms were measured with icy and rendered with coloredChromosomes. Chromosomes are aligned at the centromere and the minimum complement (n) only is shown. Species are ranked in order of evolutionary divergence.

## Discussion

We have assembled four new whole chloroplast genomes (Miscanthidium capense, Miscanthidium junceum, Miscanthus x giganteus, Narenga porphyrocoma and Sarga versicolor and we have assembled low copy number genomic loci from Sarga versicolor, Miscanthus saccharuflorus, Miscanthus x giganteus, Miscanthus transmorrisonensis and Miscanthus sinensis cv. Andante, Narenga porphyrocoma as well as Miscanthidium junceum, Saccharum hybrid LCP85-384, Saccharum spontaneum SES245A and Saccharum hybrid RB7234. These data have been integrated with previously published low copy number gene data (Fig. 2) and previously published chloroplast data (Fig. 3 and Fig. 4) to yield the most comprehensive analysis of the origins of Saccharum s.l. performed to date.

MinION sequencing of the Narenga porphyrocoma chloroplast yielded two isoforms with the SSC region inverted between them. Both isoforms were present at almost equal concentrations. This finding agrees with recent analyses of plant chloroplast and indicates that chloroplasts in plants exist as two isoforms (Walker et al. 2015; Turudić et al. 2021) (Fig. 6).

**Figure 6.**
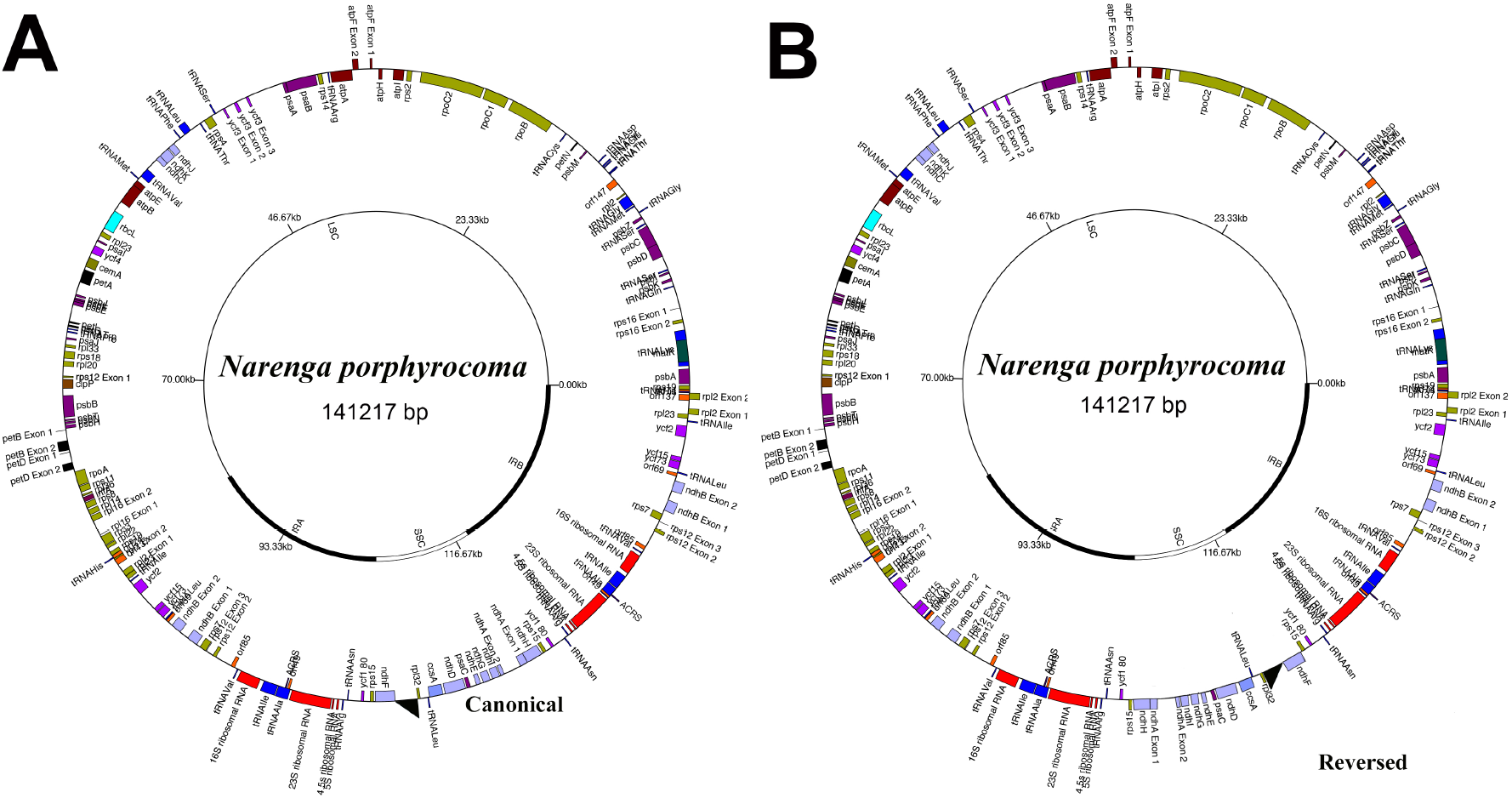
Schematic images for two chloroplast isoforms of *Narenga porphyrocoma*. Shown are schematic drawings for the two chloroplast genomes isoforms of *Narenga porphyrocoma* annotated in the present study. Left (A) is the canonical form with the SSC in forward orientation and right (B) is the form with the SSC inverted. The inner track shows the extent of the large single copy region (LSC), small single copy region (SSC) and the two inverted repeats (IRA and IRB). Images were drawn with GenomeVx prior to finishing with Adobe Illustrator.

Both low copy number gene and whole chloroplast phylogenetics clearly reveal that genus *Erianthus* is not monophyletic, with the Old World Species (*Erianthus* sect. *Ripidium*) clearly being evolutionarily very divergent from *Saccharum*. This supports the placement of these former *Erianthus* species not within *Saccharum*, but within the genus *Tripidium* (the type species for which is *Tripidium ravennae*).

The New World Erianthus species (now placed within *Saccharum* and represented by *Saccharum villosum, Saccharum asperum, Saccharum giganteum* and *Saccharum angustifolium*) along with *Miscanthidium* are clearly allopolyploids, with an A and B genome. Comparison of the gene and chloroplast phylogenies reveals that the A genome (the female parent) was derived from a *Saccharum s.s*. ancestor. *Miscanthidium* and *Erianthus* are monophyletic and divergent from each other, though they share a common ancestry. *Saccharum narenga* has a different ancestry, sharing an A genome with *Miscanthidium*, though its B genome derives from a more recent sister genome to *Saccharum* (which is supported by the chloroplast phylogeny).

*Saccharum* s.s. is monophyletic and shows no allopolyploidization outside its own genus (modern sugarcane hybrids are a complex of three genomes: *Saccharum spontaneum, Saccharum officinarum* and *Saccharum cultum* [Lloyd Evans and Joshi (2016)]). Though we could recover the *Saccharum spontaneum* copy of the low copy number genes within the *Saccharum* hybrid LCP85-384 genome; despite exhaustive gene assembly attempts we could not find any other divergent forms of the low copy number genes. This rather gives the lie to the ‘saccharum complex’ theory, demonstrating that sugarcane has not hybridized outside the boundaries of *Saccharum s.s*. and is not a melange of different progenitor species.

*Miscanthus* is also monophyletic and, again, shows no evidence of allopolyploidization outside its own genus. *Pseudosorghum fasiculaire* sits as an outgroup to *Miscanthus*, and its genome shows no evidence of secondary gene copies due to ancestral hybridization. As such, allopolyploidization within *Miscanthus* seems to have happened early in the evolutionary history of *Miscanthus* itself. It seems that the early lineages leading to *Miscanthus sinensis* and *Miscanthus sacchariflorus* hybridized reciprocally with each other, resulting in a chromosomal fusion event, and it was these hybrids that led to modern *M. sinensis* and *M. sacchariflorus* lineages.

The low copy number gene phylogeny indicates that the B genome of *Miscanthidium* and *Erianthus* diverged first. An ancestral form of this lineage hybridized with an early form of the ‘A’ genome lineage and it was this hybrid that led to *Miscanthidium* and *Erianthus*. *Miscanthidium* remained in Africa, whilst *Erianthus* species traversed Asia to reach the Americas by means of the Beringia land bridge (Zimov et al. 2012).

In *Narenga*, only two gene copies were found, one corresponding to the A genome and one that serves as an outgroup to *Saccharum s.s*. This was confirmed by transcript capture and MinION sequencing. The *Sacharum s.s*. outgroup being the female lineage, as demonstrated by the chloroplast phylogeny. We do not know from our data if *Narenga* started as a B-genome A-genome hybrid and was originally part of the *Miscanthidium* clade (but this seems likely).

Mapping chromosome numbers onto the divergence backbone (Fig. 7) we see that the majority of the lineages have different chromosome numbers. We start with *Sarga* (n=5) which makes it likely that the A genome was also n=5. There was a genome duplication held in common between *Miscanthus* and *Saccharum* (Kim et al. 2014), so the common ancestor of both these lineages had n=10 chromosomes. We would expect (simplistically) that *Miscanthidium* and *Erianthus* would have n=5 + n = 10 (ie n=15) as the base chromosome number. Indeed, studies show that the base chromosome numbers in both these groups is n=15.

**Figure 7.**
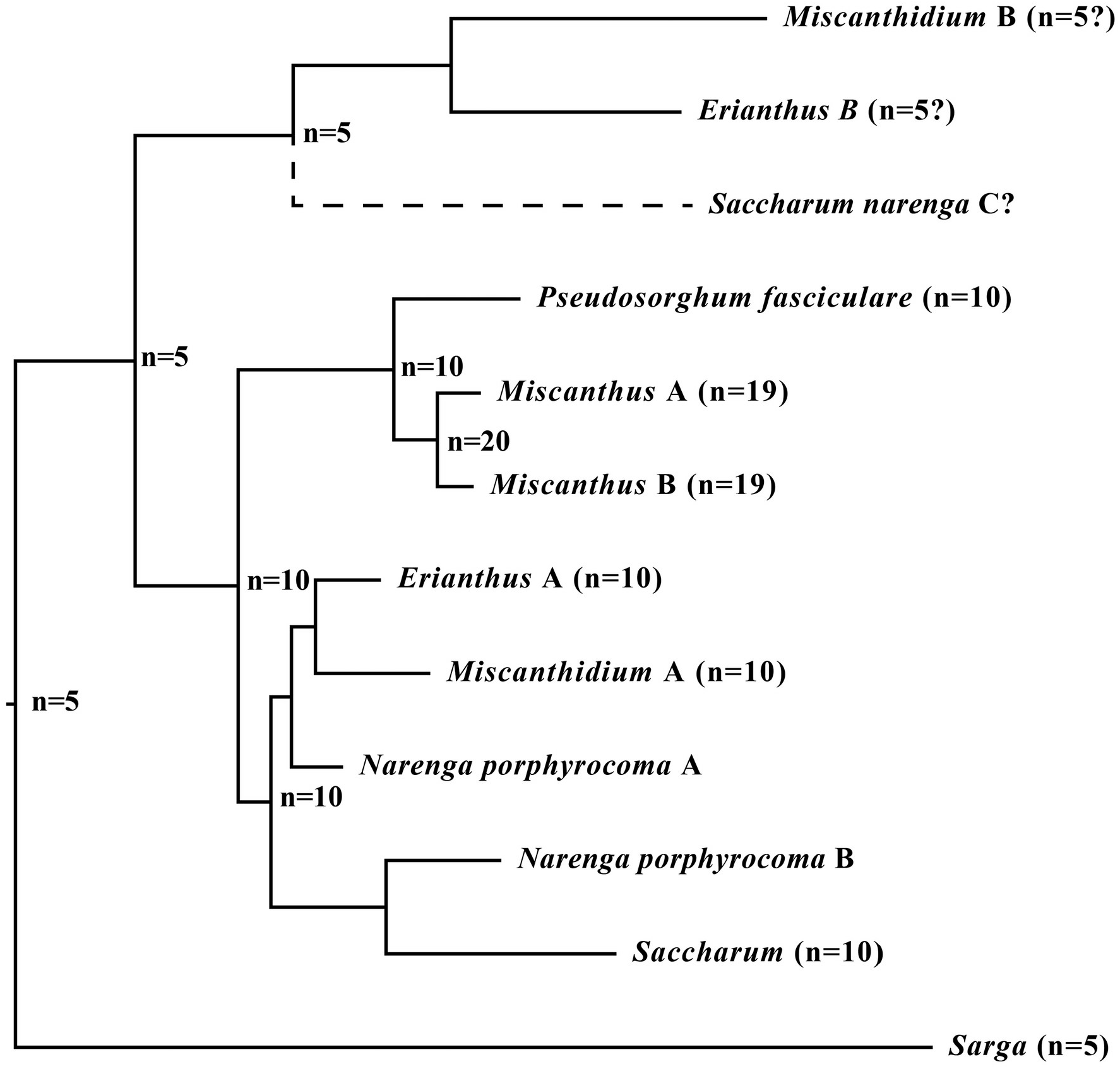
Schematic of chromosome number evolution in the Saccharinae. Shown is a schematic dendrogram representing a possible route of chromosomal evolution within the core Saccharinae, attempting to explain why different base chromosomal counts are observed within different genera within the subtribe. Numbers at nodes give starting chromosome counts before divergence, numbers at the tips give the species level chromosome counts.

The base chromosome number in *Pseudosorghum* is 10, but in Miscanthus it’s 19. We predict that there was a chromosome doubling in *Miscanthus* due to allopolyploidization between ancestral *Miscanthus* lineages, but two chromosomes fused, leading to n=19 as the base chromosome number (this leads to the satellite chromosome seen in *M. sinensis* chromosome 1 in Fig. 5). The base chromosome number in *Saccharum* is 10, though there were probably chromosomal fusions in *S. spontaneum* giving a base number of 8 (and yielding two long chromosomes). The outlier to this nice series is *Narenga*. If this genome started as a hybrid of A and B genomes, we would expect a base number of n=15. If it’s genome derived only from the A genome then we would have a base number of n=10 (this evolutionary series is mapped schematically in Fig. 7).

Given that allopolyploidization took place in *Narenga* to an n=10 *Saccharum s.s*. ancestor, we would expect either n=25 or n=20 as base chromosomal numbers. But the base chromosome number in *Narenga* is n=15. How can these studies be reconciled? Part of the answer may come from karyotype analyses (Fig. 5). *Miscanthidium capense* and *Saccharum giganteum* (*Erianthus giganteus*), both with n=15 have small chromosomes (Hoshino and Davidse, 1988; Burner and Webster, 1994). However, *Narenga* has a mix of larger and smaller chromosomes, as well as a single chromosome with a clear satellite, indicating that some of the chromosomes resulted from chromosomal fusion. Thus, in *Narenga*, a combination of chromosome loss and fusion could have led to the reduction of n=20 or n=25 (probably the most likely) to n=15. As a result, base chromosome numbers in *Sarga*, *Pseudosorghum*, *Miscanthus* and *Saccharum* are different. *Miscanthidium*, *Erianthus* and *Narenga* have the same base chromosome numbers n=15, but *Miscanthidium* and *Erianthus* have small chromosomes whilst *Narenga* has a mix of short and long chromosomes. Moreover, *Miscanthidium* has a distribution in Africa, whilst *Erianthus* species are found throughout the Americas and probably reached the continent via Beringia, as did many other grass species (Zimov et al. 2012). *Narenga* represents a species with an Asian distribution (from India to China) but with a rump population in Africa (Ethiopia), indicating that this species might have arisen in Africa and colonized Asia via a land bridge at the base of Arabia during a global sea-level minimum (Bailey et al. 2007).

## Conclusion

For the first time, we have assembled and annotated the complete chloroplast genome of two Miscanthidium species. Integrating our previous transcript-based assembly of the *N. porphyrocoma* chloroplast with the newly-assembled chloroplasts of *Miscanthidium junceum*, *M. capense, Miscanthus x giganteus* and *Sarga timorense* along with enriched low copy number datasets we present comprehensive phylogenies of the Saccharinae. We demonstrate that *Miscanthidium* and *Erianthus* (excluding *Tripidium* species) are monophyletic and separate from both *Saccharum* and *Miscanthus* (indeed, they are allopolyploid, whilst *Miscanthus* and *Saccharum* only show hybridization within their own genus).

*Miscanthidium* and *Erianthus* are also monophyletic, though they are sister taxa. We therefore support the use of *Miscanthidium* for the African species *Miscanthidium junceum* (formerly *Miscanthus junceus*) and *Miscanthidium capense* (formerly *Miscanthus capensis* syn *Miscanthus ecklonii*). *Erianthus* contains the type species for *Erianthus* (*Erianthus giganteus*, syn *Saccharum giganteum*) and our data supports the use of *Erianthus* for this grouping, which is distinct from *Saccharum* in terms of evolutionary origin and chromosome numbers). Moving *Miscanthus ecklonii* from *Miscanthus* into *Miscanthidium* means that *Miscanthus* loses its type species. However, as Hitchcock also defined *M. japonicus* (Trin.) Andersson (basionym *Saccharum floridulum* Labillardière; described in 1824), which is synonymous with *Miscanthus floridulus* (Labil.) as a second type for *Miscanthus*, then *M. floridulus* now becomes the type species for *Miscanthus s.s*.

*Narenga* is much more difficult to place. It has the same basic chromosomal number as *Erianthus* and *Miscanthidium*, but a different chromosome length pattern. It is a hybrid of a species ancestral to *Miscanthidium* and *Erianthus* with a *Saccharum* ancestor (the rump population for this species in Ethiopia indicate it may have originated in Africa [http://www.kew.org/data/grasses-db/www/imp09051.htm]). Thus, it could be placed in *Miscanthidium* or in *Saccharum* (chloroplast analyses would indicate a placement in *Saccharum*). However, its different origins, chromosomal fusions in its ancestry, base chromosome number of 15 and physical characteristics place it outside *Saccharum s.s*. We therefore conclude that it should be within its own genus and that its preferred name should be *Narenga porphyrocoma* (Hance ex Trimen) Bor.

It should also be noted that, in terms of the Saccharum complex, which is an interbreeding population, *Narenga porphyrocoma* is the closest genus to Saccharum, followed by *Miscanthidium* and *Erianthus*, *Pseudosorghum* and *Miscanthus*. As only these species lie within the about 3.4 to 4.2 million year window for hybridization in the wild (Lloyd Evans and Joshi 2016), this represents the effective limit of the Saccharum complex. Within a broader sense, both gene and chloroplast phylogenies show that all the above genera should be included in the Saccharinae, along with *Sarga*. Gene-based phylogenies indicate that *Microstegium*, *Sorghastrum* and *Polytrias* might also be included in the Saccharinae.

Chloroplast phylogeny indicates the inclusion of *Sorghum*, *Imperata* and *Pogonatherum*. However, gene-based and chloroplast based phylogenies disagree on the placement of *Sorghum* and *Pogonatherum*. We do not have equivalent samplings on the other taxa. As a result more work needs to be done to confirm the true extent of the Saccharinae, though we demonstrate that current definitions of both the *Saccharum* complex and the Saccharinae subtribe are too broad and require revision.

Our findings cast doubt on the validity of the ‘Saccharum complex’ as a group of interbreeding genera that led to the evolution of modern sugarcane. Indeed, we find no evidence for the introgression of any species outside *Saccharum* into the genome of sugarcane. Moreover, only four genera: *Pseudosorghum*, *Miscanthus*, *Miscanthidium* and *Narenga* lie within the 3.8 million year window in which wild hybridization is possible within the *Saccharinae* and thus these genera circumscribe the extent of the Saccharum complex.

## Supporting information

Capture primers for Narenga transcripts

PCR amplification primers for chloroplasts

List of chloroplast genomes used in phylogeny

## Author Contributions

DLlE and SVJ conceived the initial experiment, DLlE extended and amended the experiment design. SVJ provided the SASRI plant materials. DLlE designed PCR primers, performed the sequence assemblies, alignment and phylogenetics, developed all software scripts, analysed and interpreted the data and wrote the paper. BH captured and sequenced the Narenga porphyrocoma chloroplast and low copy number transcripts. SVJ and BH proofread the paper. All authors read and agreed to the final version of the manuscript.

## Acknowledgements

We thank Mr E. Albertse for performing the DNA isolation, PCR amplifications and DNA extractions for *Miscanthidium capense* and Mr K-L Mah for collecting the *Narenga porphyrocoma* leaf samples. All sequences from this study have been deposited in EMBL/GenBank. Assemblies performed from third party data and all alignments and phylogenetic trees have been deposited in Dryad.

## Supplementary Materials

All datasets from this study, including sequence assemblies from third party SRA datasets and optimized alignments have been deposited in the Dryad repository: . Alignments and phylogenies have been deposited in TreeBase: TB2:S21154 and TB2:S21155. Sequence assemblies were deposited in EMBL/GenBank under the project identifiers: PRJEB17861 and PRJEB22229. Supplementary Appendices 1, 2 and 3 are available with the on-line version of the journal.

