## Supplementary material for "Comparative whole plastome and low copy number phylogenetics of the core Saccharinae and Sorghinae": List of chloroplast genomes used in phylogeny

| Species | Voucher or Accession | GenBank Accession | Reference |
| --- | --- | --- | --- |
| *Arundinella deppeana* | XAL:Clark et al. 1680 | KU291490.1 | Burke et al. 2016 |
| *Arthraxon prionodes* | PI<ITA>:659331 | KU291471.1 | Burke et al. 2016 |
| *Chrysopogon serrulatus* |  | KU961864.1 | Welker et al. 2016 |
| *Eriochrysis laxa* |  | KU961863.1 | Welker et al. 2016 |
| *Eriochrysis villosa* |  | KU961860.1 | Welker et al. 2016 |
| *Saccharum spontaneum* | SES205A | LN896360.1 | Lloyd Evans and Joshi 2016 |
| *Saccharum spontaneum* | SES234B | LN849912.1 | Lloyd Evans, D. (submitter) |
| *Saccharum hybrid cultivar* | Q165 | LN896359.1 | Lloyd Evans and Joshi 2016 |
| *Saccharum hybrid cultivar* | SP80-3280 | AE009947.2 | Calsa Jr et al. Calsa jr et al. 2004 |
| *Saccharum hybrid cultivar* | NCo310 | AP006714.1 | Asano et al. 2004 |
| *Saccharum officinarum* | IJ76-514 | LN849913.1 | Lloyd Evans and Joshi 2016 |
| *Miscanthus sacchariflorus* | Hercules | LN869218.1 | Lloyd Evans, D. (submitter) |
| *Miscanthus sinensis* | Andante | LM735682 | Lloyd Evans, D. (submitter) |
| *Miscanthus floridulus* | PI295762 | LN869215.1 | Lloyd Evans, D. (submitter) |
| *Sarga timorense* |  | KF998272.1 | Kepers et al. (submitter) |
| *Sorghum bicolor* | BTx623 | EF115542.1 | Saski et al. 2007 |
| *Imperata cylindrica* | DEK:Burke 21 | KU291466.1 | Burke et al. 2016 |
| *Pogonatherum paniceum* |  | KU961859.1 | Welker et al. 2016 |
| *Eulalia aurea* | PI<ITA>:12153 | KU291499.1 | Burke et al. 2016 |
| *Sorghastrum nutans* | DEK:Wysocki s.n. | KU291482.1 | Burke et al. 2016 |
| *Hyparrhenia subplumosa* | PI<ITA>:12665 | KU291500.1 | Burke et al. 2016 |
| *Diheteropogon amplectens* | var. catangensis voucher PI<ITA>:12585 | KU291497.1 | Burke et al. 2016 |
| *Themeda sp.* | Saarela 1833 | KU291484.1 | Burke et al. 2016 |
| *Iseilema macratherum* | PI<ITA>:257760 | KU291468.1 | Burke et al. 2016 |
| *Capillipedium venustum* | PI<ITA>:11713 | KU291493.1 | Burke et al. 2016 |
| *Bothriochloa alta* | DEK:Duvall s.n. | KU291492.1 | Burke et al. 2016 |
| Tripidium ravennae | PRJEB20532 | PRJEB20532 | Lloyd Evans, D. (submitter) |
| *Tripidium arundinaceum* | SA-E1 | PRJEB20532 | Lloyd Evans, D. (submitter) |
| Tripidium arundinaceum | OK76-57 | PRJEB20532 | Lloyd Evans, D. (submitter) |
| Tripidium kanashiroi |  | PRJEB20532 | Lloyd Evans, D. (submitter) |
| *Ischaemum afrum* | PI<ITA>:364924 | KU291467.1 | Burke et al. 2016 |
| *Rottboellia cochinchinensis* | ISC<USA-IA>:Clark et al. 1698 | KU291481.1 | Burke et al. 2016 |
| *Coix lacryma-jobi* | M. Duvall s.n. 26May 2006 (DEK) | FJ261955.1 | Leseberg and Duvall 2009 |
| *Zea mays* | B73 | AY928077.1 | Schnable et al. 2009 |
| *Zea luxurians* |  | KR873424.1 | Orton 2015 |

Supplementary Material 2

GenBank accessions along with voucher accessions and associated publications (if available) for all chloroplasts employed for phylogenetic analyses in this study.
